# What matters to a mouse? Effects of internal and external context on male vocal response to female squeaks

**DOI:** 10.1101/2024.10.15.618481

**Authors:** Lauren R Brunner, Laura M Hurley

## Abstract

House mice adjust their signaling behavior depending on the social context of an interaction, but which aspects of context elicit the strongest responses from these individuals is often difficult to determine. To explore how internal and external contextual factors influence how of house mice produce and respond to social signals, we assessed how dominant and subordinate male mice differed in their ultrasonic vocalization (USV) production in response to playback of broadband vocalizations (BBVs, or squeaks) when given limited access to a stimulus female. We used a repeated measures design in which each male was exposed to two types of trials with different odor conditions: either just female odors (Fem condition) or female odors in addition to the odors of potential competitors, other males (Fem+Male condition). The presence of odors from other males in this assay served as a proxy for an “audience” as the male interacted with the stimulus female. These conditions were replicated for two distinct cohorts of individuals: males exposed to the odor of familiar competitors in the Fem+Male condition (Familiar odor cohort), and males exposed to the odor of unfamiliar competitors in the Fem+Male condition (Unfamiliar odor cohort). By assessing dominance status of the focal individual and familiarity of the “audience”, we are able to explore how these factors may affect one another as males respond to BBVs. Dominants and subordinates did not differ in their baseline vocal production (vocalizations produced prior to squeak playback) or response to squeaks. However, all groups, regardless of dominance status or odor condition, reduced their vocal production in response to BBV playback. The presence of unfamiliar male odor prompted mice to decrease their baseline level of calling and decrease the complexity of their vocal repertoire compared to trials that only included female odor, and this effect also did not differ across dominance status. Importantly, the presence of male odor did not affect vocal behavior when the male odor was familiar to the focal individual. These findings suggest that mice alter their vocal behavior during courtship interactions in response to cues that indicate the presence of potential competitors, and this response is modulated by the familiarity of these competitor cues.

## Introduction

The ability to process and respond appropriately to conspecific signals is crucial for animals as they navigate social interactions [1–3]. How an individual processes and responds to signals may heavily depend on the context of the situation [4–6], and the role of context in vocal communication in particular is highly documented across taxa [7–11]. Research on how contextual influences such as the presence of competitors and potential mates, seasonality, changes in steroid hormones, and resource abundance affect song production and song composition in birds has revealed that many aspects of communication depend on context, and prompted advances and questions on how context can influence the signaling in other species [12,12–16]. These include house mice, studied not only as a model of rodent communication signals, but also for their potential applicability to human communication disorders [17,18].

House mice (*Mus musculus*) produce a variety of vocalizations that can vary in usage, structure, and meaning across different contexts [1,10,11,19,20]. Vocal communication in mice is prominent during courtship interactions, in which calls are used to communicate receptivity and sexual motivation [1,10,21–24]. One type of call, ultrasonic vocalizations (USVs), are high-frequency calls (30-110 kHz) produced by both males and females during social interactions [1,20,24–26]. These calls are produced in large numbers by males during opposite-sex interactions, and are often referred to as “courtship calls” because of their prosocial function to facilitate mating [1,24,27]. Male USV production thus may indicate male courtship effort and motivation [22]. Mouse vocal repertoires, or the composition of different types of syllables individuals use, are complex and can vary depending on factors such as age, genetics, and behavioral situation [10,28–30]. One type of USV in particular, 50 kHz harmonic USVs, tend to be longer in duration and more complex structurally than other classes of USVs. These calls are closely associated temporally with incidents of male mounting behavior [10].

Another type of call, broadband vocalizations (BBVs), or squeaks, are lower-frequency calls with a broader frequency range than USVs, featuring many harmonics in. Both male and female mice produce BBVs across varying contexts, including during distress, handling, and conspecific interactions [26,31–33]. When produced by females during the early stages of courtship interactions, these calls are closely correlated with females performing physical rejection behaviors like kicking, lunging, and attempted escape [11,34]. Thus, during courtship, female-produced BBVs may be negatively valenced, and function to express a lack of female receptivity to sexual advances by males. In accordance with the connection of squeaks to physical rejection, male mice reduce their USV output in response to BBV playback during limited interactions with female mice [35]. However, little is known about what sorts of contextual factors may influence this response to BBVs, and which levels of context may have the most substantial impacts on mouse behavior as they interact.

The goal of this study was to explore the effects of both internal and external contextual factors on male vocal production and vocal repertoire in response to female squeaks. By examining both influences inside and outside of the focal individual, we can gain a better understanding of how these forces interact to affect behavior as a whole. As an internal contextual factor, we evaluated dominance status in all of our individuals. Some calls are more associated with the production of behaviors classified as “more dominant”, such as aggression and courting [20]. Dominant and subordinate mice tend to differ in their vocal responses to female cues, though the direction of this difference varies across studies. For example, one study found that dominant male mice call more quickly and more often compared to subordinate males in response to female stimuli [36], while in contrast, another found that subordinates called more than dominants when interacting with a female [21]. Thus, the effects of dominance status on vocal production in male mice appear to be highly-context dependent and worthy of further investigation.

In addition to the playback of BBVs, the external contextual factor of interest to us was the presence of non-vocal cues that may alter male response to BBVs: specifically, odor from other male mice. Audience effects, in which individuals adjust their behavior in the presence of additional individuals, are well-documented examples of context-dependent communication [37–39]. Studies in songbirds and domesticated fowl have revealed that the presence of audiences during courtship signaling can influence not only the acoustic properties of song and song repertoire, but also the motivation to interact with their partner [40–43]. Previous studies have established that certain strains of mice are susceptible to audience effects when responding to female cues [37,44]. For example, Seagraves et al. (2016) found that increases in vocal production in response to live and anesthetized male audience members may be due to signalers shifting their vocalizations to be directed to males, potentially in a competitive display. Thus, the presence of an audience may cause mice to adjust their vocal output and the function of their calling may change, similar to the strategic changes influenced by audiences across other contexts [37,43,45–47]. On the other hand, competitive male odor cues can decrease male vocal behavior and courtship effort when interacting with a female [21,48]. Thus, audience effects can vary depending on the context of an interaction.

Importantly, defining what constitutes an “audience” and an “audience effect” has been debated across the literature since the term’s conception [38,39]. For example, some argue that true audience member(s) do not engage in the exchange of signals with those in the communication interaction [39,49]. Some theorists and studies support that audience effects may not be limited to the physical presence of an individual acting as an audience, and that cues that indicate the proximity of a “potential” audience may be sufficient to promote these changes in signaling behavior [50]. For example, male newts (*Lissotriton boscai*) decrease their courtship effort when the area in which they are interacting with a female contains chemical cues from other males, potentially indicating a “chemically-mediated” audience effect [48]. Similarly, subordinate male mice that were evaluated for vocal production during an interaction with a female called less if they were tested in the same cage in which a dominant male and female had previously interacted, compared to if they were tested in a cage with no social odors [21]. Thus, the researchers concluded that odor cues left behind by the urine of dominant male were sufficient to affect subordinate male vocal behavior during an interaction with a female, which aligns with the framework of audience effects. Because odor cues can imply audience presence without the possibility of active signal exchange, investigating audience effects elicited by these cues can ensure that the audience cannot take part in the signaling interaction, thus the role of the audience as a “non-targeted receiver” is maintained [39,50].

In order to assess the effects of a simulated audience, in the form of conspecific male odor cues, and dominance status on male behavioral response to BBVs, we employed a recently established paradigm [35] that allows for the decoupling of BBVs and their frequently associated physical rejection behaviors. We predicted that the presence of male odor during these interactions would prompt males to decrease their vocal output in response to BBV playback more so than if no male “audience” was present, and that this effect would be dependent on dominance status. It has been reported that dominant mice are more likely to adjust their behavior in response to social challenges than subordinate mice [51–53], so we predicted that dominant males would alter their behavior in response to a simulated social competitor more than subordinates by showing a larger decrease in USV production during BBV playback if male odor was present.

We investigated the role of familiarity of the perceived “audience” in this study by assessing behavior of male mice in two distinct cohorts: one exposed to unfamiliar male conspecific odor, and the other exposed to odors from a male cage-mate with which they had an established dominance relationship. Familiarity was of particular interest to us because identity and status of audience members can alter behavior during signaling [54]. We predicted that more robust differences between dominants and subordinates would be seen in the familiar odor cohort because of the established dominance relationship between the signaler and the audience member.

Including cues that may indicate an audience, allows us to greater understand how males are responding to vocal courtship rejection signals (BBVs) in a more naturalistic environment, where they are likely to be confronted with the odors of competing males [55,56]. By assessing both dominance status and familiarity of the audience to the focal individual, we can further parse apart what role identity plays in this interaction. Surprisingly, we found that dominance status did not affect vocal behavior, while the presence of additional male odor dampened vocal production, but only when the odor came from a male unfamiliar to the focal individual.

## Methods

### Animals

This study was carried out in strict accordance with the recommendations in the Guide for the Care and Use of Laboratory Animals of the National Institutes of Health. The protocol was approved by the Bloomington Institutional Animal Care and Use Committee at Indiana University (Protocol number: 21–020). Focal subjects consisted of seven-week-old male CBA/J mice (N = 23) (The Jackson Laboratory, Bar Harbor, ME). Mice were housed in a 14:10 light:dark cycle with food and water *ad libitum*. All focal and stimulus mice were housed in same-sex pairs from at least one week prior to behavioral assays until the end of the experiments in order to allow for the establishment and maintenance of a social dominance relationship within the cage. Dominance status in male mice was evaluated using multiple measurements of a standardized tube test in which cage-mates were simultaneously released at opposite ends of a clear plastic tube and allowed to interact until only one mouse remained in the tube [35,57]. The mouse that retreated from the tube was classified as subordinate, while the mouse that remained in the tube longer was classified as dominant. The tube used was narrow enough to prevent the mice from passing over one another [35,50]. All housing pairs underwent 3-4 tube tests with no more than one test per day, and with tests occurring no more than 4 days prior to the trial dates. Almost all mice were consistent in their dominance status across all of their tests, aside from two pairings that switched dominance status after the first test, but were consistent for the final three tests. At least two days prior to experimental trials, all mice were given opposite-sex experience with novel female CBA/J mice (The Jackson Laboratory, Bar Harbor, ME) through multiple 10-minute interactions across the span of 2-3 days [11]. These interactions occurred in clean cages, in which each focal male was allowed to interact alone with a stimulus female. All mice received approximately the same amount of social experience prior to trials (50-80 minutes total with an average of 67.14 minutes, SD = 14.85). A student’s *t*-test confirmed that individuals across the two cohorts received statistically similar amounts of social experience (*t*(26) = −0.142, p = 0.888). All stimulus animals for experimental trials were sexually experienced adult female CBA/J house mice novel to the focal mouse.

### Behavioral assay and experimental conditions

A split-cage playback paradigm previously established in our lab [35] was employed to evaluate the effects of vocal playback on male mouse vocal behavior. In this paradigm, the focal mouse and stimulus female are placed on opposite sides of a 12 inch x 6 inch x 6 inch cage with a plexiglass barrier in the middle with a small hole at the bottom of the barrier that allows for limited nose-to-nose social investigation. The presence of a live female, opposed to only female odor or an anesthetized female, is necessary to promote consistent robust vocalization from the focal male during this assay [35]. Approximately 30g of dirty bedding containing urine, feces, and body odor [58,59] from the stimulus female and her cage mate was placed on the focal male’s side of the cage to promote robust vocalizations [35,36], while clean bedding was placed on the female’s side of the cage. Dirty bedding from female cages was obtained after 24-48 hours after the bedding was placed with the animals. A microphone (CM16/CMPA, Avisoft Bioacoustics, Glienicke/Nordbahn, Germany) and Canon VIXIA video camera were placed above the cage on the focal male’s side to allow for recording of ultrasonic vocalizations (Fig 1A) and BBV playback (Fig 1B), and non-vocal behavior, respectively, during the interaction. The entire cage and recording setup were located in a sound-proof chamber (IAC Acoustics, Naperville, IL) in a room separate from the area in which the colony was housed. All trials were performed between 9:00 AM – 4:00 PM during the light portion of the animal’s light cycle. Mice were habituated to the sound chambers 1 day prior to their experimental trials for 60 minutes. On the day of the trials, the focal male mouse was placed in the experimental cage for 60 seconds, or until they started producing USVs, and subsequently, the focal female was then placed on the opposite side of the cage. Mice were in this cage for 15-minute playback trials that consisted of 5 minutes of silence (a baseline period), followed by 5 minutes of exemplar BBV playback through an ultrasonic speaker powered by Avisoft next to the cage, followed by another five minutes of silence (a recovery period). Hence forth, these time periods will be referred to as test periods. The segment of BBV playback was created using five repeated 1-minute segments of BBVs collected from a previous direct interaction between a male and a female during a period of frequent physical rejection and BBV vocalization [35]. Each call during this period was replaced with a single exemplar BBV with a duration, peak frequency, and percentage of non-linear segments within one standard deviation of the mean for all BBVs in the interaction in order to eliminate possible differences in behavior over the course of the playback due to variation in BBV structure [35]. The BBV segment was played at a loudness calibrated to be approximately matched to the intensity that male mice would experience from BBVs during natural interactions with females, with a maximum of 104.9 dB [35].

**Fig 1:**
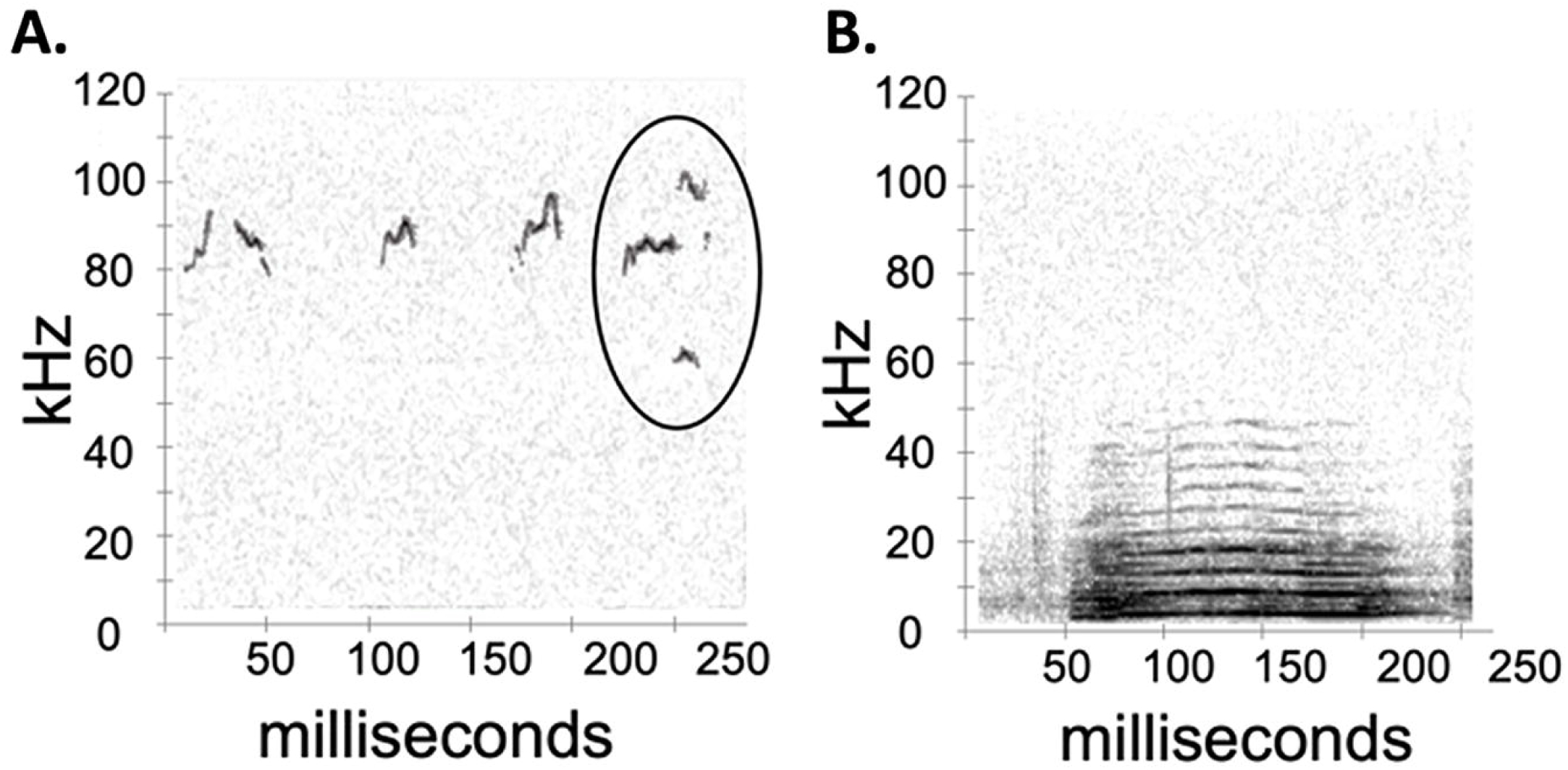
Representative spectrograms of (A) Ultrasonic vocalizations (USVs) and (B) Broadband vocalizations (BBVs). The circled vocalization in (A) is an example of a 50 kHz

Previous use of this paradigm in our lab has shown that males decrease their USV production during BBV playback relative to the baseline and increase back to baseline levels during the recovery. This response to BBV playback when males are physically separated from the stimulus female indicates that playback of BBVs can alter male courtship vocal behavior even when the correlated physical rejection behaviors are absent [35]. For the current study, a modified version of this paradigm was created to allow for two distinct experimental conditions that altered the available olfactory cues: a female-scent-only condition in which only bedding from the stimulus female’s home cage was placed on the focal animal’s side of the cage (Fem condition) and a female scent + male scent condition in which the focal male was exposed to the 30g of bedding from the stimulus female as well as 30g of bedding from a cage of male mice (Fem+Male condition; Fig 2).

**Fig 2:**
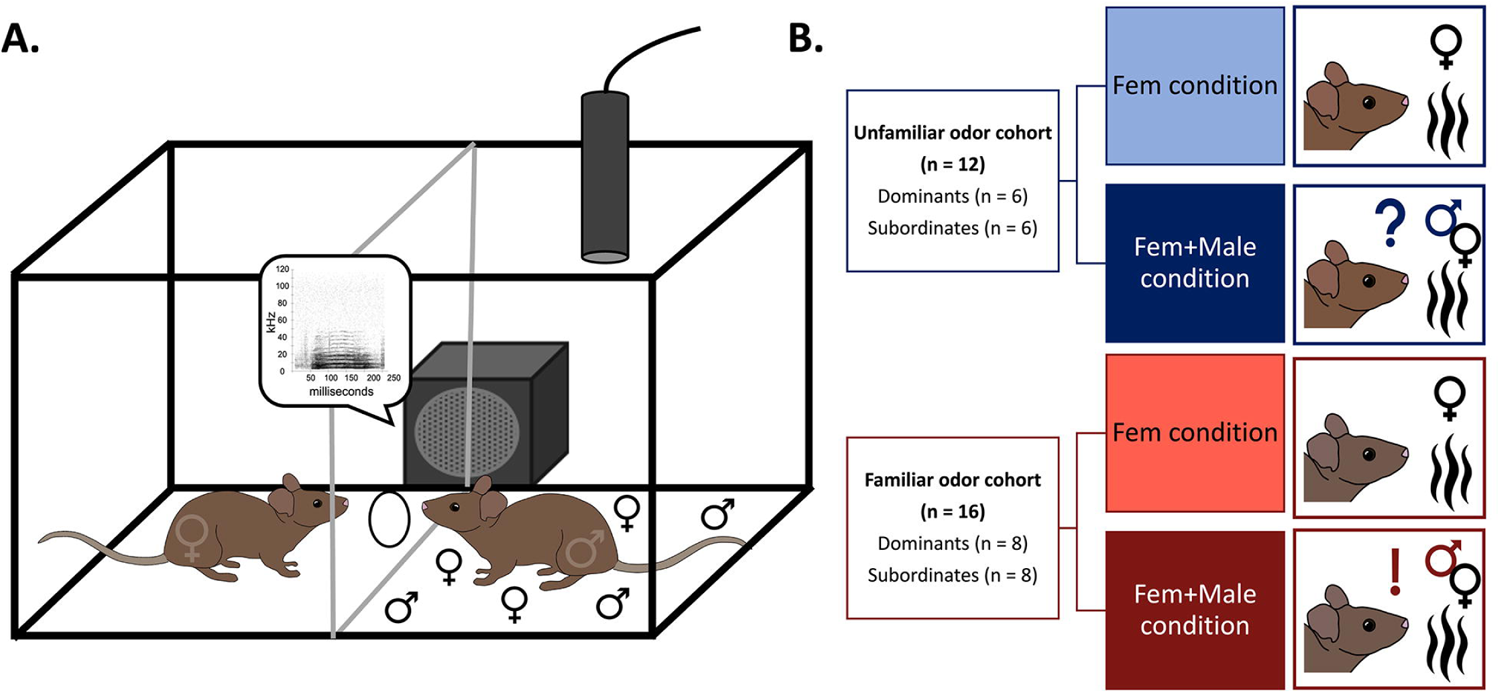
Experimental assay and design. (A) Split-cage paradigm used to evaluate the effects of only BBV playback on male vocal behavior, modified to create the Fem+Male condition for this study. Male and female symbols present on the male mouse’s side of the cage represent the presence of dirty bedding from male and female mice. (B) Experimental groups broken up by familiarity cohort, dominance status, and odor condition. A question mark icon is used to illustrate unfamiliarity with the odor stimulus in the trial, and an exclamation point icon is used to illustrate familiarity with the odor stimulus in the trial.

Both assay conditions were applied to the two separate experiments with separate cohorts of animals described in the following sections. These experiments differed based on the origin of the male bedding used in the Fem+Male condition, described below, while the Fem condition trials in both experiments were identical in design. All individuals in both experiments/cohorts underwent both a Fem condition trial and a Fem+Male condition trial to account for individual differences in vocal behavior. Different trials for the same mice occurred approximately 24 hours apart and the order of the trials was counterbalanced across all individuals.

### Experimental design

Experiment 1 - Unfamiliar male odor cohort: In the Fem+Male condition, individuals in the unfamiliar male odor cohort (n=12) were exposed to equal amounts of dirty bedding from the stimulus female’s cage and dirty bedding that came from a cage of males unfamiliar to them. These “odor donations” were from the other cages of males used in this experiment. No males from separate cages ever interacted with one another, thus all of the “odor donations” came from males novel to the focal individuals. Dirty bedding containing urine, feces, and body odor from unfamiliar male odor donors was collected after the unfamiliar donor male was allowed to explore the cage for 2 hrs. The Fem+Male odor condition in this experiment was designed this way to simulate the presence of an unfamiliar male competitor during the limited interaction with the female.

Experiment 2 - Familiar male odor cohort: In a subsequent experiment, a new cohort (n=16) was established as the familiar male odor cohort. For these individuals, in the Fem+Male condition they were exposed to equal amounts of dirty bedding from the stimulus female’s cage and dirty bedding from their own home cage, in order to simulate the presence of their familiar cage mate: the other half of their dominance pairing. Dirty bedding containing urine, feces, and body odor from the familiar male odor donors was collected from a cage that held the focal male and their cage mate for 2 hrs.

### Audio analysis

USVs from all of the interactions were counted individually by trained individuals using Avisoft SasLab Pro software (Avisoft Bioacoustics, Glienicke/Nordbahn, Germany). A total of 74,859 USVs were counted manually during the course of this study. Durations of the USVs were automatically calculated by Avisoft. In order to assess specific components of the individuals’ vocal repertoires, all USVs were manually categorized as either non-harmonic or 50 kHz harmonic USVs (Fig 1). 50 kHz harmonics were defined as syllables with at least one harmonic band at 40-50 kHz, as per Hanson and Hurley’s (2012) criteria [10], while all other USVs were categorized as non-harmonic if they did not possess this specific frequency band (Fig 1). 50 kHz harmonic USVs were specifically denoted because they are highly correlated to male mouse mounting behavior and could potentially be used to promote receptivity in females during courtship interactions [10,35].

### Non-vocal behavior analysis

Non-vocal behavior was analyzed using Behavioral Observation Research Interactive Software (BORIS) [60]. The odor condition and cohort of the focal individual was blinded to the analyst for each trial. Because male and female investigative behavior may be related in this assay [35], the duration of time both the focal male and stimulus female spent at the hole in the plexiglass divider was scored for all individuals. Investigation of the window was defined as the mouse placing their nose at the entrance of (< 1 cm away from the entrance) or through the opening in the barrier. Investigation of holes in a barrier that separates a focal and stimulus mouse has also been historically scored to understand overall investigative behavior in assays like these [7,35].

### Statistical analysis

Statistical analyses were performed in IBM SPSS Statistics 28. A Pearson correlation was used to evaluate the relationship between the baseline USVs produced in the Fem condition and the baseline USVs produced in the Fem+Male condition across all individuals to assess consistency in baseline vocal production. Four Pearson correlations were used to evaluate the relationship between the investigative behavior of the focal male and the stimulus female in each cohort, for each odor condition. Due to the difference in the Fem+Male conditions across the two cohorts, separate general linear models were used for each cohort for analysis of vocal and non-vocal behavior. For all models that used repeated measures, the mouse’s individual ID served as the repeated measure.

To assess the effects of BBV playback across the 5-minute test periods (baseline, playback, and recovery) and odor treatment on number of non-harmonic calls, a three-way repeated-measures ANOVA was used for the unfamiliar odor cohort, and an identical three-way repeated-measures ANOVA was used for the familiar odor cohort (within-subjects factors: test period and odor treatment; between-subjects factor: dominance status. Analyses on the effects of odor condition and dominance status on baseline number of USVs in each interaction and the duration of baseline USVs produced were conducted using identical two-way mixed ANOVAs for the unfamiliar odor cohort and familiar odor cohort (within-subjects factor: odor treatment; between-subjects factor: dominance status). Interactions among all three factors were included in these models.

In order to investigate how the composition of vocal repertoires responded to our manipulations, we characterized the proportion of harmonic USVs out of all USVs in each test period of the interaction. To assess the effects of BBV playback across test periods and odor treatment on proportion of harmonic calls produced, proportions of harmonic calls across test periods were calculated as: number of harmonic calls in test period/number all calls in test period. These proportions were then compared using identical binary logistic regressions for each cohort in which each vocalization was coded as either harmonic or non-harmonic [61–63] (within-subjects factors: test period and odor treatment; between-subjects factor: dominance status). Binary logistic regressions were also used to evaluate the effects of odor condition and dominance status on the baseline proportion of harmonic USVs for the unfamiliar odor cohort and familiar odor cohort (within-subjects factor: odor treatment; between-subjects factor: dominance status).

To assess the effects of BBV playback across test period and odor treatment on non-vocal behavior, two identical three-way repeated-measures ANOVAs were used, one for each cohort (within-subjects factors: test period and odor treatment; between-subjects factor: dominance status). There was an error with one individual’s video file for the last ten minutes of the trial, so this file was excluded from analysis for these measures. Interactions among all three factors were included in these models.

The following datasets violated the ANOVA assumption of normality and thus were normalized using square root transformations in order to run analyses of variance: USVs across test periods in the unfamiliar odor cohort, and female investigative behavior in the unfamiliar odor cohort. Small violations of the homogeneity of variance requirement were noted, but as these models were the best fit aside from this fact, and because ANOVAs are robust to small violations of this assumption when sample sizes are equal, we proceeded with ANOVAs [64]. Benjamini-Hochberg corrections (Q value = 0.05) were conducted in order to account for multiple comparisons [65], and Bonferroni tests were performed as posthoc analyses.

The possibility of outliers in the datasets was investigated using analyses of the number of standard deviations that each value deviated from the mean of the dataset, and double-checked using Tukey’s method [66,67]. No outliers were revealed in any of the datasets.

## Results

### Individual baseline vocal production in both odor conditions is positively correlated

All individuals underwent both a Fem condition trial, in which focal individuals were only exposed to the odor of the female stimulus animal, and a Fem+Male condition trial, in which focal individuals were exposed to both odor from the female stimulus and odor from an additional male conspecific (Fig 2B). A Pearson Correlation revealed that the baseline vocal production across both trials was strongly positively correlated across individuals in both the familiar odor cohort and unfamiliar odor cohort (r = 0.735, p < 0.001).

### Dominance status does not alter behavior across cohorts or odor conditions

Dominance relationships for each cage consisted of a dominant individual and a subordinate individual, and the placement of individuals within these dominance relationships remained stable throughout multiple rounds of tube tests. Out of 14 dominance pairings, only 2 pairings changed status at all throughout the tube tests. Both of these switches occurred during the second test and the pairings were subsequently stable throughout the rest of the three tests. When incorporating dominance status in our statistical models as a between-subjects factor, there were no significant effects of dominance status in any behavioral measures (Tables 1-3).

**Table 1.** Output of models assessing the effects of BBV playback, Fem or Fem+Male odor condition, and dominance status on total USV production and harmonic USV production in both cohorts.

|  |  | Number of USVs in Unfamiliar odor cohort <sup>a</sup> |  | Proportion of 50 kHz Harmonics in Unfamiliar odor cohort <sup>b</sup> |  |
| --- | --- | --- | --- | --- | --- |
| *Effect* | *df* | *F* | *p* | *F* | *p* |
| Test Period | 2 | 50.538 | <.001 | 4.925 | **0.011** |
| Odor Condition | 1 | 12.026 | **0.006** | 8.429 | **0.005** |
| Dominance Status | 1 | 0.019 | 0.892 | 0.702 | 0.406 |
| Test Period x Odor Condition | 2 | 0.223 | 0.802 | 2.236 | 0.116 |
| Period x Dominance Status | 2 | 0.614 | 0.551 | 0.131 | 0.877 |
| Odor Condition x Dominance Status | 1 | 0.742 | 0.409 | 0.022 | 0.882 |
| Odor Condition x Test Period x Dominance Status | 2 | 1.203 | 0.321 | 1.093 | 0.342 |

<sup>a</sup> Nonharmonic USVs were assessed with three-way mixed ANOVAs.
<sup>b</sup> Proportion of 50 kHz harmonic USVs were assessed using binary logistic regressions.

### Additional male odor during limited social interactions decreases baseline vocal output depending on the familiarity of male odor

The presence of unfamiliar male odor during the interaction (Fem+Male condition) caused significantly lower baseline levels of USVs, USVs during the first 5 minutes of the interaction, compared to when only female odor was present (Fem condition) (Two-way mixed ANOVA: F_1,10_ = 10.005, p = 0.01). The presence of familiar male odor however produced no significant difference in baseline USV levels compared to when only female odor was present (Two-way mixed ANOVA: F_1,14_ = 0.084, p = 0.776) (Fig 3A) (Table 2).

**Figure 3:**
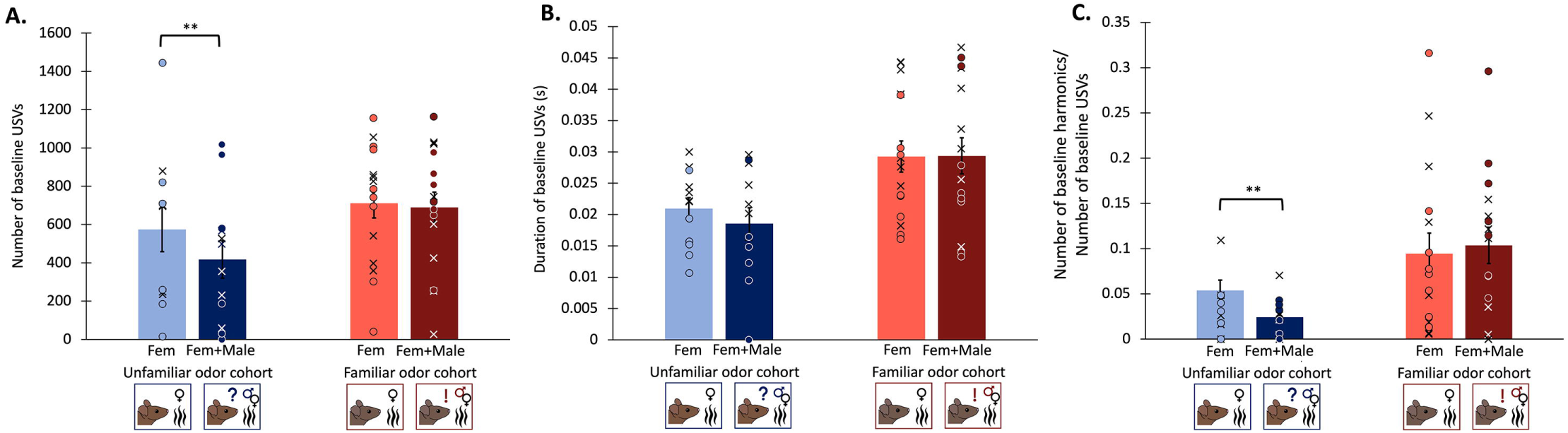
Properties of vocal repertoires. (A) Baseline USVs produced in the Fem condition vs the Fem+Male condition. (B) Duration of baseline USVs in seconds. (C) Baseline proportion of harmonic USVs (# baseline harmonics/# baseline USVs). Proportion of baseline harmonic USVs was calculated as baseline harmonic USVs /(baseline harmonic USVs + baseline nonharmonic USVs). Blues represent the unfamiliar odor cohort, reds represent the familiar odor cohort. Bars represent the means of the experimental groups, and means from dominants and subordinates are pooled due to the lack of effect of social status. Error bars are calculated from standard errors. Individual points of data noted with dominants as X’s

**Table 2.** Output of two-way mixed ANOVAs assessing the effects of Fem or Fem+Male odor condition and dominance status on baseline USV production, baseline harmonic USV production, and duration of baseline USVs in both cohorts.

|  |  | Baseline number of USVs in Unfamiliar odor cohort |  | Baseline 50 kHz Harmonics in Unfamiliar odor cohort |  | Duration of baseline USVs in Unfamiliar odor cohort |  |
| --- | --- | --- | --- | --- | --- | --- | --- |
| *Effect* | *df* | *F* | *p* | *F* | *p* | *F* | *p* |
| Odor Condition | 1 | 10.005 | **0.01** | 10.752 | **0.004** | 5.285 | 0.044 |
| Dominance Status | 1 | 0.04 | 0.846 | 0.036 | 0.852 | 0.051 | 0.826 |
| Odor Condition x Dominance Status | 1 | 0.919 | 0.36 | 2.101 | 0.163 | 0.576 | 0.465 |

\* p value is not statistically significant after correcting for multiple comparisons.

**Table 3.**
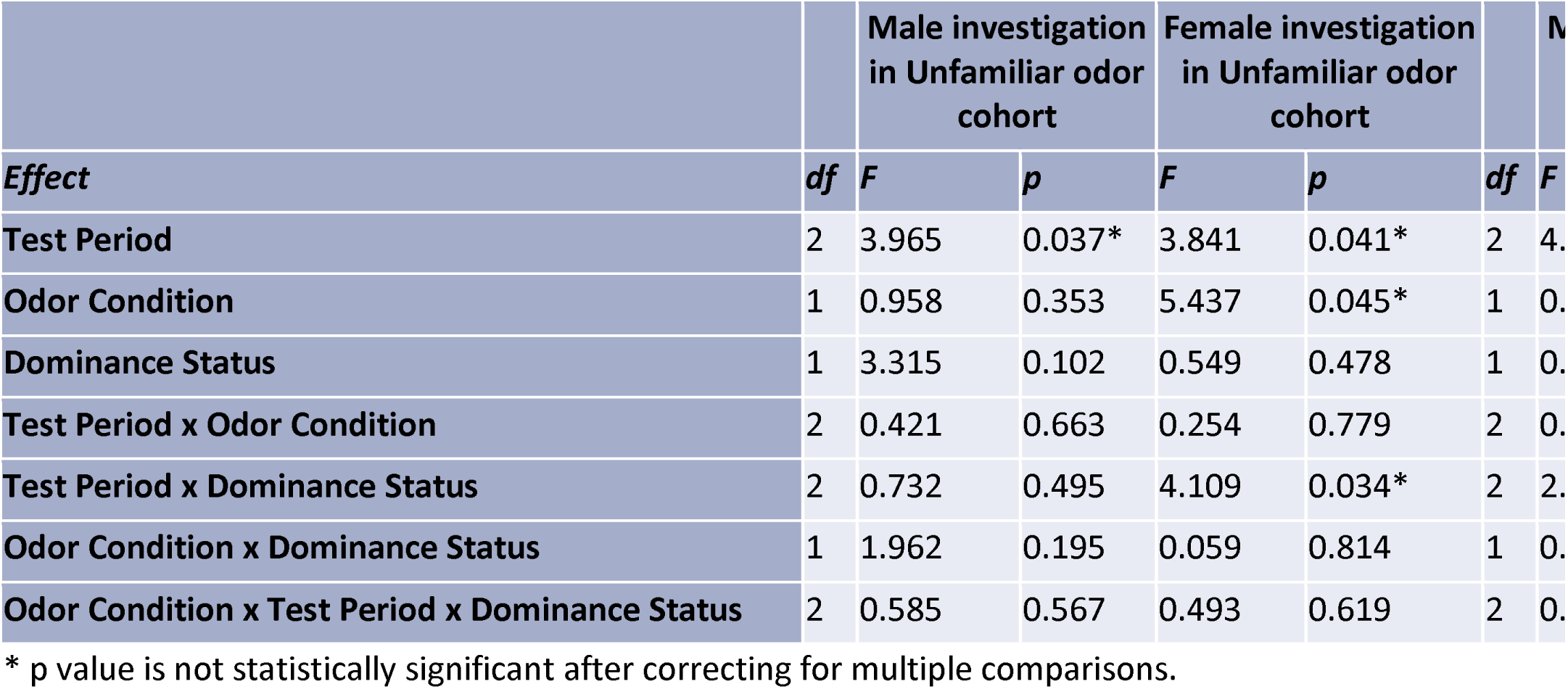
Output of three-way mixed ANOVAs assessing the effects of BBV playback, Fem or Fem+Male odor condition, and dominance status on male and female investigation of the divider window in both cohorts.

| *Effect* | *df* | Male investigation<br>in Unfamiliar odor<br>cohort |  | Female investigation<br>in Unfamiliar odor<br>cohort |  | *df* | *F* |
| --- | --- | --- | --- | --- | --- | --- | --- |
|  |  | *F* | *p* | *F* | *p* |  |  |
| Test Period | 2 | 3.965 | 0.037* | 3.841 | 0.041* | 2 | 4. |
| Odor Condition | 1 | 0.958 | 0.353 | 5.437 | 0.045* | 1 | 0. |
| Dominance Status | 1 | 3.315 | 0.102 | 0.549 | 0.478 | 1 | 0. |
| Test Period x Odor Condition | 2 | 0.421 | 0.663 | 0.254 | 0.779 | 2 | 0. |
| Test Period x Dominance Status | 2 | 0.732 | 0.495 | 4.109 | 0.034* | 2 | 2. |
| Odor Condition x Dominance Status | 1 | 1.962 | 0.195 | 0.059 | 0.814 | 1 | 0. |
| Odor Condition x Test Period x Dominance Status | 2 | 0.585 | 0.567 | 0.493 | 0.619 | 2 | 0. |

\* p value is not statistically significant after correcting for multiple comparisons.

Because USVs can vary widely in duration [1], we measured the duration of all baseline USVs (both harmonic and nonharmonic). A trend of males producing longer baseline USVs in the Fem condition compared to the Fem+Male condition in unfamiliar male cohort was noted (Two-way mixed ANOVA: F_1,10_ = 5.285, p = 0.044), though this result was not significant after correcting for multiple comparisons (Table 2). No effects of odor condition on duration of baseline USVs were observed in the familiar male odor cohort (Two-way mixed ANOVA: F_1,14_ = 0.005, p = 0.945) (Fig 3B).

### Additional male odor during limited social interactions decreases baseline usage of 50 kHz harmonic USVs depending on the familiarity of male odor

50 kHz harmonic USVs are calls highly correlated to mounting behavior in male mice [11]. In this study, harmonic USVs made up a small proportion of all calls; on average, harmonic USVs made up 7.31% (SD = 0.0701) of all USVs in the interaction. In order to assess whether the frequency of harmonic USV usage differed across conditions, for each individual we evaluated the proportions of harmonic USVs produced in each test period out of the total number of all USVs in that test period. The presence of unfamiliar male odor during the interaction (Fem+Male condition) caused mice to use a significantly lower proportion of 50 kHz harmonic USVs during the baseline period compared to when only female odor was present (Fem condition) (Binary logistic regression: F_1,10_ = 10.752, p = 0.004), though this effect was not present in the familiar male odor cohort (Binary logistic regression: F_1,14_ = 0.035 p = 0.853) (Fig 3C) (Table 2).

### Male mice decrease USV production in response to BBV playback across all conditions

In both the unfamiliar odor cohort and the familiar odor cohort, male mice reduced their calls during playback of BBVs, and then increased calling rates after playback ceased in both Fem and Fem+Male trials (Three-way mixed ANOVAs: Significant main effect of test period F_2,20_ = 50.538, p < 0.001 and F_2,28_ = 34.635, p < 0.001 for the unfamiliar and familiar cohorts, respectively). This subsequent increase in calling after the playback period did not fully recover to levels similar to baseline, however, but rather increased to a level between that observed in the baseline and the recovery. In the unfamiliar odor cohort, individuals produced more USVs in the Fem condition compared to the Fem+Male condition (Significant main effect of odor condition: F_1,10_ = 12.026, p = 0.006) (Fig 4A), though this effect was not present in the familiar odor cohort (F_1,14_ = 0.454, p = 0.511) (Fig 4B). No interaction effects between odor and test period were present in either the unfamiliar odor cohort or the familiar odor cohort (F_2,20_ = 0.223, p = 0.802, and F_2,30_ = 0.734, p = 0.489) (Table 1).

**Figure 4:**
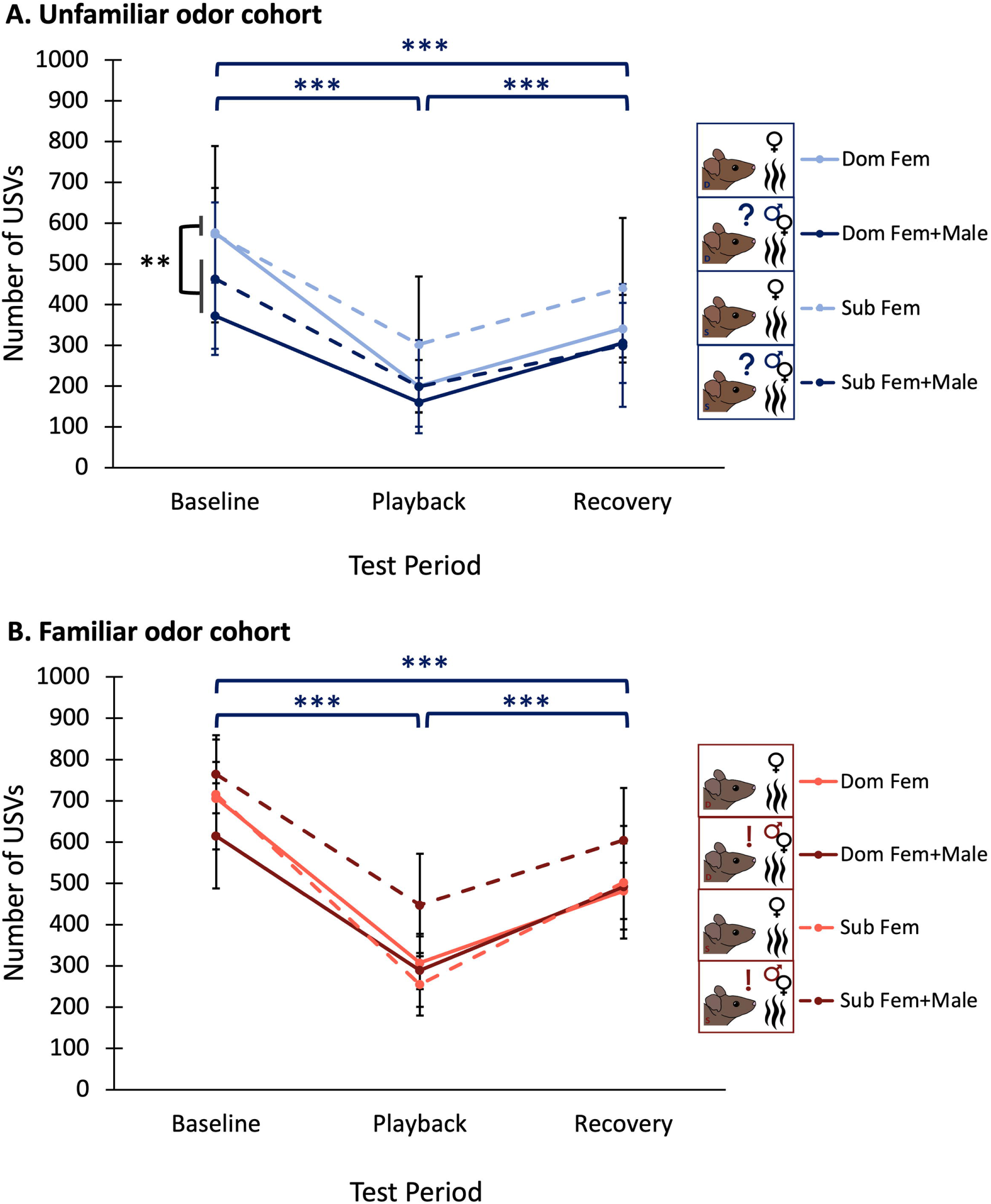
Number of USVs produced across test periods (A) in the unfamiliar odor cohort, and (B) in the familiar odor cohort. Average USV production of dominants is represented by the solid lines, and average USV production of subordinates is represented by dashed lines. Lighter lines denote average USV production of individuals in the Fem condition, while darker lines denote average USV production of individuals in the Fem+Male condition. Error bars are calculated from standard errors.

Because there was no interaction effect between test periods and any other factor in these models, our data showed that differences across odor conditions and familiarity cohorts can be attributed to differences in baseline levels of USV production, rather than differences in vocal response to BBV playback.

### Male mice increase usage of 50 kHz harmonics after BBV playback ends when exposed to unfamiliar male odor

In the unfamiliar male odor cohort, the proportion of 50 kHz harmonic calls used relative to the total number of calls produced did not change in response to BBV playback; however, after playback ended, males produced a greater proportion of harmonic calls in the recovery period (Binary logistic regression: Significant main effect of test period F_2,20_ = 4.925, p = 0.011). However, posthoc pairwise comparisons revealed this increase in harmonic usage only occurred in trials with unfamiliar male odor added (t_55_ = 2.344, p = 0.023) (Fig 5A). Overall, in the unfamiliar male odor cohort, males produced a greater proportion of harmonics in Fem trials compared to Fem+Male trials (Binary logistic regression: Significant main effect of odor condition F_1,10_ = 8.429, p = 0.005). In the familiar male odor cohort, no significant effects of any factor were observed (Table 1) (Fig 5B).

**Figure 5:**
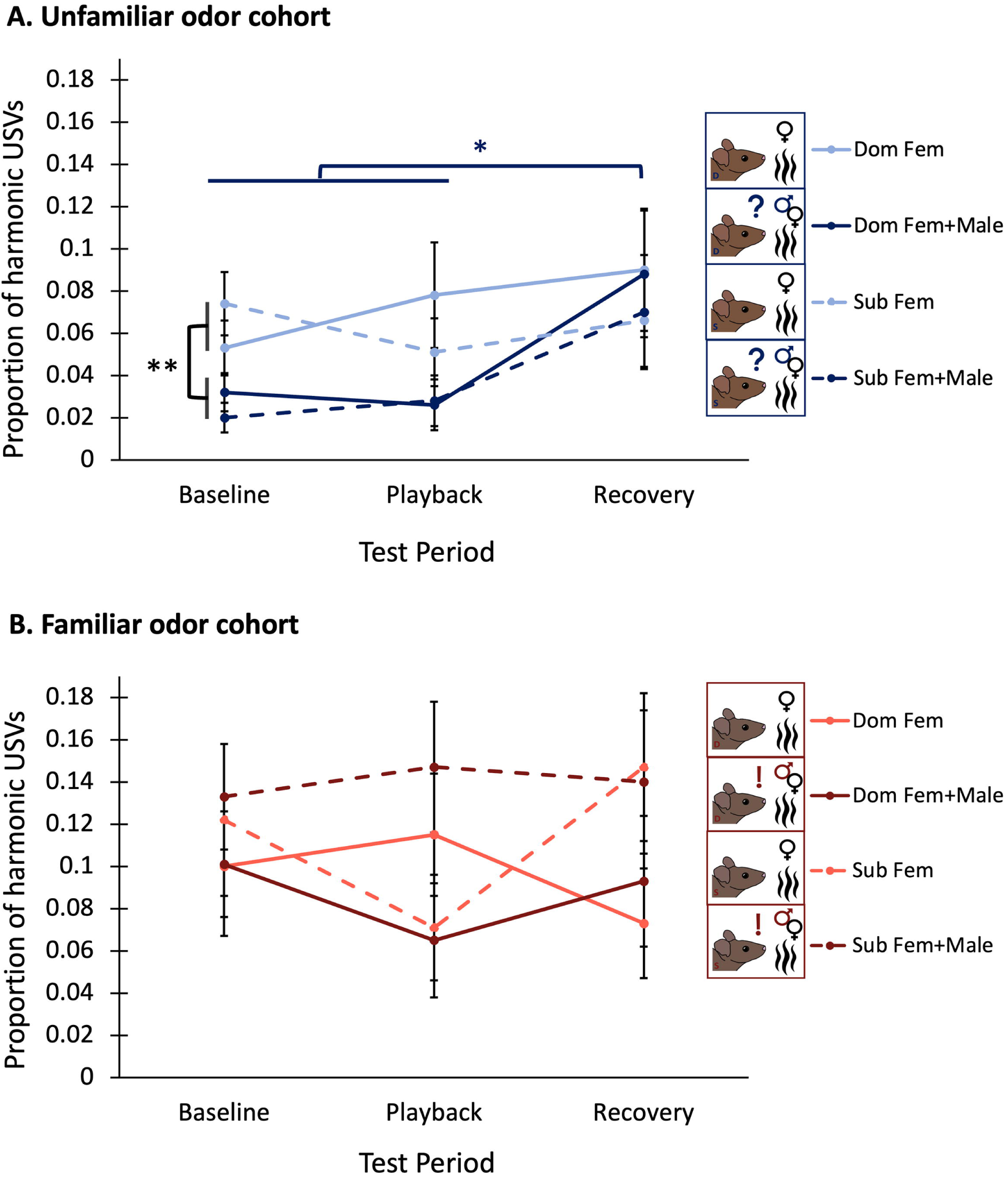
Proportion of harmonic USVs across test period out of total USVs (A) in the unfamiliar odor cohort and (B) in the familiar odor cohort. Average harmonic USV production of dominants is represented by the solid lines, and average harmonic USV production of subordinates is represented by dashed lines. Lighter lines denote average harmonic USV production of individuals in the Fem condition, while darker lines denote average harmonic USV production of individuals in the Fem+Male condition. Proportion of harmonic USVs for each test period was calculated as harmonic USVs in test period/(harmonic USVs + nonharmonic USVs in test period). The significance bar in 5A) showing increased usage of proportion of harmonic USVs in the recovery period compared to the baseline and playback period is dark blue, representing that it only applies to the Fem+Male conditions, not the Fem conditions. Error bars are calculated from standard

### Male and female investigative behavior does not significantly differ across conditions, cohorts, nor test periods

In the unfamiliar odor cohort, male investigation of the divider window appeared to slightly decrease during BBV playback, and investigative behavior did not recover after BBV playback, with no effects of dominance status or odor condition (Three-way mixed ANOVA: F_2,18_ = 3.965, p = 0.037) (Fig 6A). In contrast, in the familiar odor cohort, male investigation of the divider window appeared to slightly decrease during BBV playback and subsequently recover after playback, with no effects of dominance status or odor condition (Three-way mixed ANOVA: F_2,28_ = 4.391, p = 0.022) (Fig 6C). However, neither of these results was significant after correcting for multiple comparisons.

**Figure 6:**
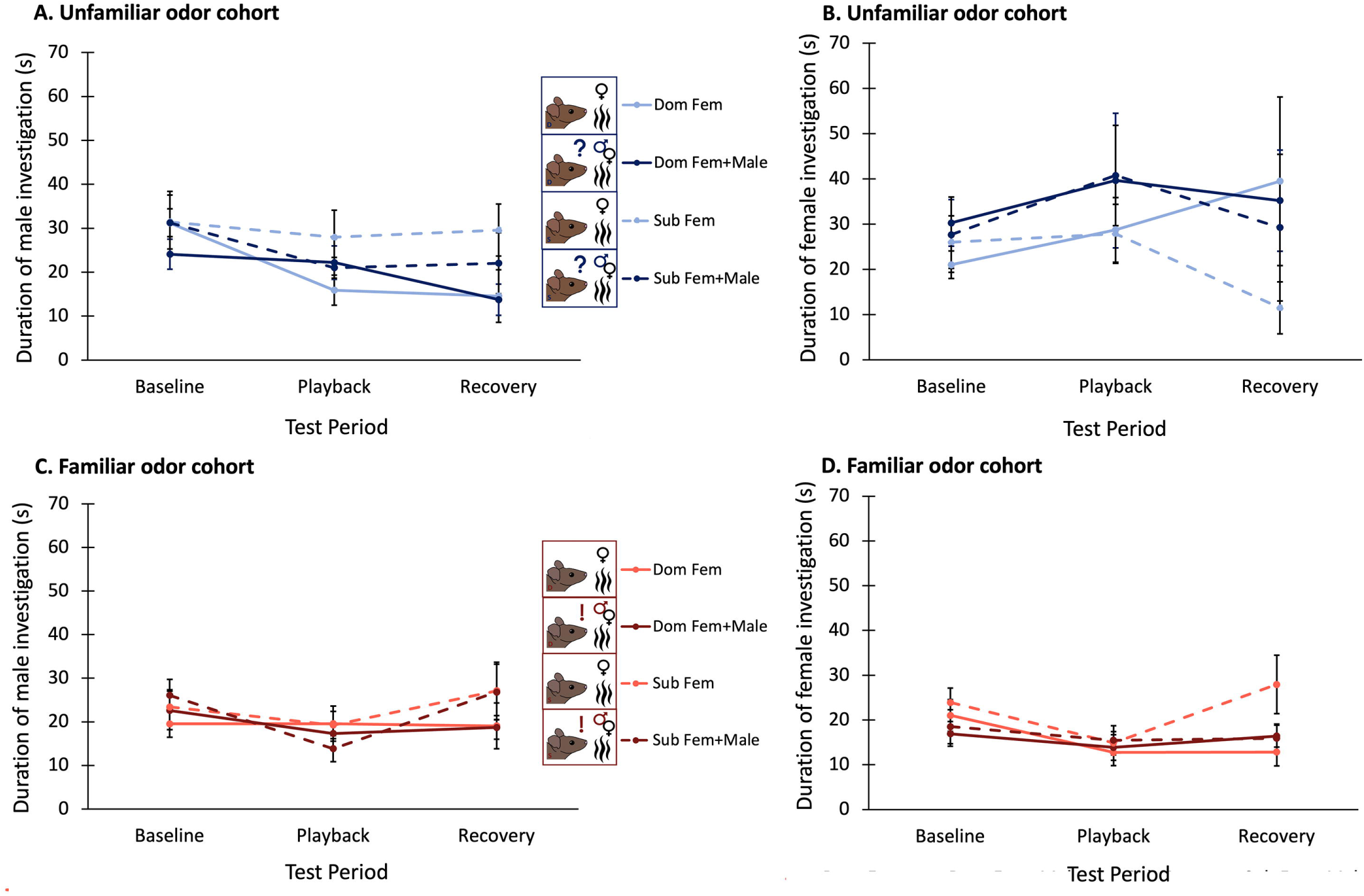
Duration of time the focal male (A and C) or the stimulus female (B and D) spent investigating the divider window across test periods in the unfamiliar odor cohort (A and B) and in the familiar odor cohort (C and D). Average duration of investigation in trials with dominant males is represented by the solid lines, and average duration of investigation in trials with subordinates is represented by dashed lines. Lighter lines denote average investigation during trials in the Fem condition, while darker lines denote average investigation in the Fem+Male condition. Error bars are calculated from standard errors.

Stimulus females paired with males in the unfamiliar odor cohort tended to investigate the divider window more during the Fem+Male condition trials (F_1,9_ = 5.437, p = 0.045), and this investigation actually increased during BBV playback, and decreased back to baseline levels after playback ended (F_2,18_ = 3.841, p = 0.041) (Fig 6B). This effect of playback was dependent on dominance status in that female investigative behavior was only affected by playback when females were paired with subordinate males, and there was no effect when they were paired with dominant males (F_2,18_ = 4.109, p = 0.034). When stimulus females were paired with males in the familiar odor cohort, they decreased their investigative behavior during BBV playback, and this behavior increased back to baseline levels when playback ended (F_2,28_ = 3.949, p = 0.031) (Fig 6D), with no effect of odor condition or dominance status. However, again, none of these results were significant after correcting for multiple comparisons.

### Additional male odor alters the relationship between male and female investigative behavior

In the Fem condition trials, the total duration of male investigative behavior did not correlate to the total duration of female investigation behaviors in either the unfamiliar odor cohort or the familiar odor cohort (Pearson correlation: r = −0.373, p = 0.259; r = 0.235, p = 0.381 respectively). In the Fem+Male condition trials, the total duration of male investigative behaviors negatively correlated to the total duration of female investigative behaviors for those in the unfamiliar odor cohort (Pearson correlation r = −0.625, p = 0.04), though this effect was not significant after correcting for multiple comparisons. In contrast, in the Fem+Male odor condition trials conducted with the familiar odor cohort, the total duration of male investigation behaviors positively correlated to the total duration of female investigative behaviors (Pearson correlation r = 0.839 p < 0.001), and this effect remained significant after correcting for multiple comparisons.

## Discussion

Behavioral decisions during courtship are highly influenced by the interplay between environmental cues, internal factors, and signals from communicating conspecifics [4]. Thus, by evaluating how individuals respond to signals in various contexts, we can begin to decipher how much weight a given factor may have on such decisions. In this study, when interactions included odor cues from unfamiliar, but not familiar, male conspecifics, male mice appeared to exhibit reduced courtship motivation compared to interactions when only female odors were present. This decrease in courtship motivation was suggested by shifts in multiple aspects of male courtship vocalizations: decreases in call production, call durations, and variety of calling repertoires. Male mice thus may be evaluating their environment for potential competitors, discerning the identity of these competitors, and scaling their courtship activities in response to these cues. The interplay between signals not only from courtship partners but also from competitors as well as social recognition of those signals is instrumental in motivating behavioral decisions during mating interactions.

### Unfamiliar male odor during courtship decreases male courtship motivation

In many taxa, vocal signaling can often reflect motivational state [68–73]. USV production is a well-established proxy for courtship motivation in male mice, as these calls function to encourage female sexual receptivity and mating [18,21,22,24,74,75]. Furthermore, out of the possible measurements for courtship motivation in mice, these signals are particularly important as they can convey information about male fitness, identity, and reproductive state, which can in turn influence female mate choice [22,26,76]. Changes in courtship motivation can be spurred by exposure to social cues [69,70]. It is well-established that auditory communication, like birdsong, exhibits signal flexibility in response to social context [77]. For example, male cowbirds will modulate the attractiveness of their songs in response to the presence of competitors, though how they modulate their songs may depend on factors like whether or not the competitors can easily access the focal individual and if the individual can gauge the motivational state of the competitor [78,79].

We found that whether males responded to the male “audience” odor was dependent on whether or not the focal male had social contact with the odor’s source. When presented with unfamiliar male odor during the interaction, males produced fewer calls during the baseline period (Fig 3A), with a shorter average length of calls than when male odor was not present (Fig 3B). Similarly, they also shifted the composition of the different types of syllables they used, their vocal repertoire [29], to produce a lower proportion of 50 kHz harmonic calls during the baseline period when unfamiliar male odor is present (Fig 3C). These responses may indicate a decrease in courtship motivation and effort. Not only is number of calls produced indicative of motivational state, but types of calls produced can also reveal information about motivation. Across taxa, increasing vocal production through the expression of complex syllables and longer calls, like 50 kHz harmonics [10,11], is associated with greater energetic costs. Longer durations of vocalizations have also been associated with increased courtship display effort in singing mice (*Scotinomys teguina)*, and female rodents tend to prefer males that produce longer “higher-effort” songs [75,80,81]. Thus, calling less and producing fewer complex harmonic calls and shorter calls thus could indicate decreased reproductive motivation.

Olfactory cues are often crucial for defining the context of an interaction, particularly opposite-sex interactions. Therefore, individuals may adjust their behavior in response to these cues, potentially to avoid expending unnecessary time and energy engaging with individuals that are not sexually receptive [82–88]. Odor cues can be used to simulate different audiences and investigate how different identity characteristics alter behavior during sexual interactions [21,48,50]. However, a previous study from Seagraves et al. (2016) found that, when presented alone, male mouse urine and male mouse body odor (in the form of saliva, tears, and fur), did not affect how males vocally responded to female odor, leading the authors to conclude that male mouse odors were not sufficient to induce an audience effect [44]. Here, we found that unfamiliar, but not familiar, male odors from dirty bedding were able to prompt a significant decrease in vocal behavior when the males were interacting with a live female stimulus mouse. Our results mirror those of several other studies that found that male odor cues were sufficient to affect courtship behavior [21,48,89]. Our results and the results from Seagraves et al. (2016) could be conflicting due to methodological factors. Primarily, our odor stimulus contained both urine and feces, and potentially body odors from allowing the males to interact freely with the bedding that was later used as the odor stimulus. Odors produced from these different sources contain different volatile compounds [58,90], thus having urine, feces, and body odor in the bedding used in the interaction could have served as a more potent stimulus more likely to induce an affect than urine or body odor alone [58].

Also in contrast to the present results, Seagraves et al. (2016) found that two males presented with female odors vocalized more when together than when alone, leading the authors to conclude that the males were acting as an “audience” for one another that prompted the increase of vocal output [44]. Another interpretation that the authors suggest for their results, however, is that the female odor was acting as a potential audience to the male-male interaction, rather than a live male audience to a male interacting with female odor. This interpretation aligns more succinctly with stricter definitions of audience effects that specify that the audience does not directly communicate with the members of the interaction [39]. Seagraves et al. (2016) compare the vocal behavior of one live individual presented with odor to two live individuals presented with odor; it is logical that the addition of another live individual that is able to vocally interact with the focal individyal would increase calling compared to when the focal individual is simply presented with female odor. Thus, it is possible that rather than a male “audience” prompting the increase in calling, it was the presence of a live interacting individual being present at all in the interaction that prompted increases in calling. Here, in both our control condition (Fem) and our experimental condition (Fem+Male), there was always a live female present to promote vocalizations. The results here conflated with the Seagraves et al. (2016) study together may suggest that the presence of female odor during a male-male interaction increases vocal production, while the presence of male odor during a male-female interaction decreases vocal production, emphasizing the effect of context of the interaction on vocal communication.

It may be logical for males to show decreased courtship motivation when smelling other males, through the reduction of and simplification of calling behaviors, because cues that indicate the presence of potential reproductive competitors can decrease sexual motivation and induce rapid behavioral plasticity [54,91]. Males may “budget” copulatory efforts depending on factors like risk of competition [50,92–98]. Across taxa, male precopulatory courtship behaviors can be reduced as the density of competitors increases [92,96]. This could be due to adaptive changes in reproductive strategies by the males or sexual interference from the competitors [92,96].

A second cohort of individuals were exposed to the odor of their cage mate, which with they had an established dominance relationship, and interestingly, when the odor was familiar to the focal individual, their vocal behavior did not differ from when no odor was present at all. Not only was there no difference in dominant and subordinate individuals (to be discussed in more detail in subsequent sections), but number of calls, duration of calls, and repertoire of calls were not affected by the familiar competitor odor at all. This important distinction between the responses of individuals to familiar compared to unfamiliar competitor odors could have occurred due to various reasons. Mice can identify individuals they are familiar with through chemical cues in urine [99]. After frequent exposure to odor cues from conspecifics, mice can habituate to these odors, and thus have a less pronounced response to them [89,100,101]. The dear enemy hypothesis predicts that familiar competitors should elicit less of a reaction from territory owners than unfamiliar competitors, due to recognition and knowledge of the benefits and costs of defending against those particular individuals [102–104]. Our results are consistent with the dear enemy hypothesis in that the cues from familiar competitors elicit less of a response (less of a decrease in calls) than cues from unfamiliar competitors. The familiarity of the male cues present could determine whether or not males pay more attention to the male cues, or ignore them and shift focus to the female. As the dirty bedding used to simulate the familiar audience came from the home cage of the focal individual and their cage mate, it is also possible that the males in this cohort could have been affected by smelling their own odors as well. A potential way for future studies to explore if, in this context, the decrease in vocal behavior in response to unfamiliar competitor odor is paired with an increase in behaviors aimed to signal to the competitor could be to assess competitive scent marking behavior of the focal males when confronted with familiar vs unfamiliar odor [89,105,106]. If males confronted with unfamiliar odor are shifting their efforts from courting the females to focusing on competing with potential competitors, we would expect to see an increase in marking behavior is correlated with a decrease in vocal behavior.

### BBVs spur decreased courtship effort in male mice

All experimental groups in this study decreased calling in response to playback of broadband vocalizations, BBVs, and no other factors measured affected this response (Fig 4). While the function of BBVs across contexts has been largely unexplored, evidence suggests that in certain courtship contexts, BBVs serve to communicate a negative affective state of the signaler [26,34,107,108]. Surprisingly, we found that when individuals were exposed to the odor of an unfamiliar competitor, the proportion of 50 kHz harmonic USVs they used increased in the recovery period after exposure to BBV playback, and it was in this period that they used the highest proportion of harmonics in their calling, even more so than in the baseline period (Fig 5A). More in depth research will have to explore the intricacies of why the discontinuation of BBV playback might spur a rapid increase in harmonic usage only when cues from an unfamiliar potential competitor are present, though this could potentially reflect habituation to the competitor cues as the trial goes on. This result, though difficult to interpret, does however support that the cessation of BBV playback changes the trajectory of syllable usage in male mice when cues indicating the presence of an unfamiliar competitor are present.

BBVs may not always produce a negative reproductive response, and are sometimes correlated with increases in mounting and calling [11,108,109]. It is possible that the potential different functions of BBVs manifest when BBVs are paired with other nonvocal behaviors, and the isolated vocal characteristics of the BBVs may indicate a negative valence, and thus promote decreased courtship effort in receivers without access to additional cues from the female.

Our results exemplify that these signals are significant and attention-grabbing to male mice, seemingly regardless of certain identity and environmental factors. BBVs and their correlated behavioral responses remain a largely untapped line of research; and very few studies have explored the effects of BBV playback isolated from the associated physical rejection behaviors often correlated with BBV production. Greater research devoted to this area of mouse communication could help elucidate how different internal and external factors may affect responses to these signals.

### Lack of an effect of dominance status

In this study, dominance status did not significantly affect any vocal or nonvocal behavioral measurements in either the unfamiliar odor cohort or the familiar odor cohort. This was not what we predicted, as it is generally accepted that dominant male mice produce more USVs than subordinate individuals in response to a novel female [110,111], though USVs are not a standard way to measure dominance as they do not directly measure access to mates, but rather reflect mating effort [22]. Because USV production is highly dependent on other contextual factors, the relationship between USV production and dominance is likely influenced by finetuned differences in social history and situational context rather than being simply and strictly linear [112]. For example, certain studies have found the relationship between dominance and increased USV production depends on the types of cues present and the recency of social defeat, and can even reverse [7,19,36]. Interestingly, in this study, the individual with the highest number of baseline calls in all conditions (both odor cohorts in both the Fem and Fem+Male conditions) was subordinate (Fig 3A), though there was no overall statistically significant effect of dominance on any measurements. Despite observations that this strain of mice may not exhibit overt severe dominance in their hierarchies [113], all housed pairs in this experiment showed consistent and reliable dyadic dominance relationships through multiple rounds of the tube test. Thus, depending on the context of an interaction, dominance may affect behaviors such as offensiveness and aggression during tube tests, while not manifesting in indirect courtship assays like that used in this study. It is also possible that, because of the smaller sample size used in this study, decreased statistical power in the statistical models we used may make it more difficult to ascertain an effect of dominance. Future studies on fewer experimental groups with larger sample sizes within each group could resolve this issue.

Dominant and subordinate males also tend to differ in their response to social challenges [114–116]. Dominant rodents tend to be more susceptible to negative health consequences and behavioral impairment following social challenges compared to subordinate males [51,52,117,118], however this is most often exhibited during direct interactions with social competitors. In this study, dominant males and subordinate males were not differentially affected by odors of competitor males during courtship, thus odors alone may not be sufficient to induce this effect of the dominance relationship. Salience of cues, interaction context, social history, and genetic factors may all contribute to how the effects of dominance manifest during courtship [110,116,119–121], and more research is needed to tease out which of these factors are most influential on determining the effects of dominance.

## Conclusions

By looking at how these multiple factors intertwine to affect behavioral decisions during courtship, we can begin to assess what “matters” to a mouse as they navigate mating and when and where to exert effort during courtship. Future studies should incorporate larger sample sizes to increase statistical power when considering interactions between the variety of variables we investigated. This study supports the potential negative valence of a relatively ill-explored female signal: broadband vocalizations, or squeaks, in certain contexts. We determined that while males robustly adjust their behavior in response to these calls, the degree of this behavioral response was not altered by any of the internal or external factors we manipulated. Furthermore, this study posits that cues indicating the presence, or “audience”, of potential competitors can alter male mouse courtship motivation and behavior, but this effect is dependent on social history with the audience members. Our results support that certain identity factors, like dominance status, do not affect the relationship between these cues and the observed behavioral response.

## Supporting information

Data file and README for data file

## Acknowledgements

The authors would like to thank and acknowledge Dr. Kayleigh E. Hood, Ph.D. for providing mentorship, and guidance regarding the split cage assay, and Morgan Leever, M.P.A., M.S. for consulting regarding statistical analysis.

## Notes

### Competing Interest Statement

The authors have declared no competing interest.

### Summary of Updates

Introduction and discussion clarified; revised figures; supplemental data files updated

