## Supplementary material for "What matters to a mouse? Effects of internal and external context on male vocal response to female squeaks": Data file and README for data file: README.rtf

Dataset title: Behavioral Data for “What matters to a mouse? Effects of internal and external context on male vocal response to female squeaks”

Corresponding Author Information
Name: Lauren R. Brunner
Institution: Indiana University


Principal Investigator Information
Name: Laura M. Hurley
Institution: Indiana University


Date of data collection: 2019-09-02 - 2021-08-01

Geographic location of data collection: Bloomington, Indiana, USA 

DATA OVERVIEW

File: Brunner & Hurley Data.xlsx
Number of variables: 24
Number of cases/rows: 56. Each row represents the behavioral data of a single trial. As each individual underwent two trials (one of each condition), there are two rows per individual.
Variable list and variable information:
	Individual: An identifier for the individual mouse from which the data in that row originated from.
	Dominance status: Whether the mouse was denoted as dominant (D) or subordinate (S).
	Condition: Whether the data in that row corresponds to the trial with female odor only (Fem) or had added male odor in it as well (Fem+Male)
	Total USVs: The total number of ultrasonic vocalizations for the 15-minute trial.
	Average duration of Calls - Baseline: The average duration of the USVs emitted during the baseline period (first five minutes of the interaction). Measured in seconds.
	Number of USVs - Baseline: The number of USVs emitted during the first five minutes of the interaction.
	Number of USVs - Playback: The number of USVs emitted during the second five minutes of the interaction (during the BBV playback period).
	Number of USVs - Recovery: The number of USVs emitted during the last five minutes of the interaction.
	50 kHz Harmonic calls - Baseline: The number of USVs that were identified as 50 kHz harmonic calls in the first five minutes of the interaction.
	50 kHz Harmonic calls - Playback: The number of USVs that were identified as 50 kHz harmonic calls in the second five minutes of the interaction (during the BBV playback period).
	50 kHz Harmonic calls - Baseline: The number of USVs that were identified as 50 kHz harmonic calls in the last five minutes of the interaction.
	Male investigating - Baseline: The duration of time the focal male spent investigating the window of the barrier during the first five minutes of the interaction. Measured in seconds.
	Female investigating - Baseline: The duration of time the stimulus female spent investigating the window of the barrier during the first five minutes of the interaction. Measured in seconds.
	Both investigating - Baseline: The duration of time the focal male and the stimulus female spent investigating the window of the barrier simultaneously during the first five minutes of the interaction. Measured in seconds.
	Male investigating - Playback: The duration of time the focal male spent investigating the window of the barrier during the second five minutes of the interaction (during the BBV playback period).  Measured in seconds.
	Female investigating - Playback: The duration of time the stimulus female spent investigating the window of the barrier during the second five minutes of the interaction (during the BBV playback period). Measured in seconds.
	Both investigating - Playback: The duration of time the focal male and the stimulus female spent investigating the window of the barrier simultaneously during the second five minutes of the interaction (during the BBV playback period). Measured in seconds.
	Male investigating - Recovery: The duration of time the focal male spent investigating the window of the barrier during the last five minutes of the interaction. Measured in seconds.
	Female investigating - Recovery: The duration of time the stimulus female spent investigating the window of the barrier during the last five minutes of the interaction. Measured in seconds.
	Both investigating - Recovery: The duration of time the focal male and the stimulus female spent investigating the window of the barrier simultaneously during the last five minutes of the interaction. Measured in seconds.
	Male investigating - total: The duration of time the focal male spent investigating the window of the barrier during the entire interaction. Measured in seconds.
	Female investigating - total: The duration of time the stimulus female spent investigating the window of the barrier during the entire interaction. Measured in seconds.
	Both investigating - total: The duration of time the focal male and the stimulus female spent investigating the window of the barrier simultaneously during the entire interaction. Measured in seconds.
	Cohort: Whether the focal individual was in the Unfamiliar odor cohort (Unfamiliar) or Familiar odor cohort (Familiar).

ADDITIONAL INFORMATION
Blank cells for row 50 (Fem Condition trial for Individual 7398B) are due to an error with the video file during the last ten minutes of the file. This trial was thus when analyzing nonvocal behavior, this trial was excluded from analyses that used test period as a factor.
